# Large-effect loci in a duplicated genomic region modulate reproductive phenology in salmon

**DOI:** 10.1101/2024.03.30.587279

**Authors:** Patrick D. Barry, Natasha Howe, Gregory L. Owens, Diana S. Baetscher, Katie D’Amelio, Scott Vulstek, Joshua Russell, Elizabeth Lee, Andrew Barclay, Anthony J. Gharrett, David A. Tallmon, Wesley A. Larson

## Abstract

Whole genome duplication (WGD) provides evolutionary opportunities to increase biological complexity and phenotypic diversity by selective retention of duplicated genes. We identify a small genomic region in salmon that governs complex life history variation and has diversified across four WGDs to fine-tune trait expression. Within this region, three large-effect loci on three chromosomes modulate migration timing across four species - a critical trait optimizing fitness by synchronizing reproduction and environmental conditions. Life history and genome resequencing data support intraspecific evolution of polymorphisms related to early- and late-reproducing phenotypes. We demonstrate that the repeated evolution of loci underlying reproductive timing carries the signature of WGD-facilitated adaptive evolution. This adaptation is driven by the subsequent subfunctionalization of genes within the highly conserved HPG axis, which regulates vertebrate reproduction.

## Main Text

Whole genome duplication (WGD) events are common throughout eukaryote evolution and have been associated with adaptive radiations and the reduced risk of extinction (*1*). Duplicated genes can play a key role in evolutionary novelty through neo- and sub-functionalization. Subfunctionalization partitions gene function between two copies, allowing for greater specialization, while neofunctionalization allows a gene copy to evolve into a novel function while the sister copy retains the ancestral role (*2–4*). Because duplication preserves the original gene function, it offers a distinct path to adaptive phenotypic evolution compared to other mechanisms, such as de novo mutations in single genes (*5*), structural variants (*6*), and introgression (*7*).

Following duplication, duplicated genes commonly undergo rapid deterioration of function and loss under relaxed purifying selection (*3*). A gene’s ultimate fate depends on its stoichiometric constraints and whether it arose via single gene or whole genome duplication - a distinction rooted in gene dosage balance. Whereas a single gene duplication isolates and amplifies one component, often disrupting the relative amounts of gene products critical for cell function, WGD duplicates entire interaction networks simultaneously preserving balanced gene sets. Despite this immediate balance, most recently duplicated genomes must mitigate the detrimental effects of polyploidy by undergoing rediploidization (*8*, *9*). Throughout this return to diploid meiotic pairing via genome rearrangements or sequence divergence, genes involved in transcriptional regulation, signal transduction, and development are often retained at a higher frequency (*10–13*).

Salmonids provide a useful model to study the evolutionary significance of a WGD and subsequent rediploidization process. Two rounds of WGD, 1R and 2R, occurred early in the vertebrate lineage, a third round was specific to the Teleost lineage (Ts3R), and a fourth salmonid-specific duplication (Ss4R) occurred in the most recent common ancestor ∼106 million years ago (MYA) (*14, 15*). Following the Ss4R duplication, the group has diversified into >100 species across 10 genera with extensive life history variation. Pacific salmon and their close relatives have not completed rediploidization after the Ss4R; residual tetrasomic inheritance on eight chromosome arm pairs is conserved across much of Salmoninae (*16–19*). Salmon genomes are characterized by diverse post-duplication histories. This diversity has likely played a key role in the evolution of integrated phenotypes, which involve suites of traits that contribute to complex life histories, in response to varying demographic, environmental, and ecological regimes (*15*, *20*).

Most life history diversity is encoded by many alleles of small effect spread across the genome (polygenic traits) and relatively few have large effects (*21*). Previous studies on the genetic architecture underlying phenotypic variation in salmon have focused mostly on alleles of small effect, some of which occur in duplicated regions, potentially supporting the adaptive importance of WGD (*22*, *23*). However, most phenotypic variation is controlled by many loci of small effect which makes it difficult to investigate how WGD influences specific phenotypes and requires surveys of hundreds-to-thousands of genes (*24*).

Recent research has indicated that some complex life history phenotypes in salmon (*e.g.*, age-at-maturity, anadromy vs. residency, and migration timing) are largely controlled by a single or a small number of genomic regions (large-effect loci) (*25*), but those studies did not explore links between large-effect loci and WGD. Our study identifies three large-effect loci that are associated with migration timing in multiple salmon species and located on ohnologs. These large-effect loci provide an example of how the genetic architecture controlling biological variation can evolve from WGD.

### Migration timing is associated with a duplicated genomic region

Adult migration timing (‘run timing’ hereafter), the transition of maturing salmon from marine to freshwater habitats prior to spawning, has extensive impacts on salmon populations and the ecosystems they inhabit. Run timing in salmon can be remarkably consistent across years within a single population, yet incredibly variable across populations and species (*26*). Run timing is synchronized with long-term average environmental conditions to maximize individual lifetime fitness (*27*). Within-population variation in run timing buffers interannual variability in optimal return timing, while among-population variation contributes to the portfolio of life history diversity in salmon that underpins their resilience despite constantly changing environments (*28*). In addition to individual and population level effects, variation in salmon run timing has large cascading ecological impacts; it influences the spatial and temporal distribution of a large variety of species reliant on the delivery of marine-derived nutrients to freshwater and terrestrial ecosystems (*28*, *29*).

We investigated the genetic basis of run timing in four species of Pacific salmon (*Oncorhynchus sp.*) with whole genome sequencing (WGS) and amplicon sequencing. WGS collections in our study encapsulate a range of run timing phenotypes (Table S1): the midpoint of early and late runs are separated by approximately three weeks (pink salmon, *O. gorbuscha*); seven weeks (sockeye salmon, *O. nerka*); three months (chum salmon, *O. keta*); and four months (coho salmon, *O. kisutch*). In addition to differences in run timing, each collection represents different scales of spatial overlap, including samples collected from the same stream pool (pink), an inlet tributary and outlet of the same lake (coho), two habitat types (beach/stream) within a series of interconnected lakes (sockeye), and populations separated by >1500 km in the same river (chum salmon; Fig. S1).

Analysis of WGS data for the four species revealed three chromosomal regions, shared by two or more species, with high genetic divergence between early and late runs. All species were aligned to the chum salmon genome to simplify genomic comparisons (chromosome numbers refer to the chum genome (*30*)). Genetic divergence between runs was estimated with *F*_ST_ and a local score approach to identify significantly elevated regions shared by multiple species (*31*). Mean genome-wide *F*_ST_ between run timing groups ranged from 0.002 in Pick Creek sockeye salmon to 0.024 in chum salmon (Table S2). All four species were characterized by one or more localized peaks of elevated genetic divergence (Fig. 1). Three such peaks of divergence were shared by two or more species. Pink, sockeye, and chum salmon shared a peak (mean *F*_ST_ 0.495-0.573) that spanned ∼200-kb on Chromosome (Chr) 35, chum and coho shared a peak (mean *F*_ST_ 0.315 - 0.709) that spanned ∼750-kb on Chr 29, and sockeye and coho salmon shared a peak (mean *F*_ST_ 0.106 - 0.203) that spanned ∼500-kb on Chr 8 (Fig. 1, Table S2).

**Figure 1:**
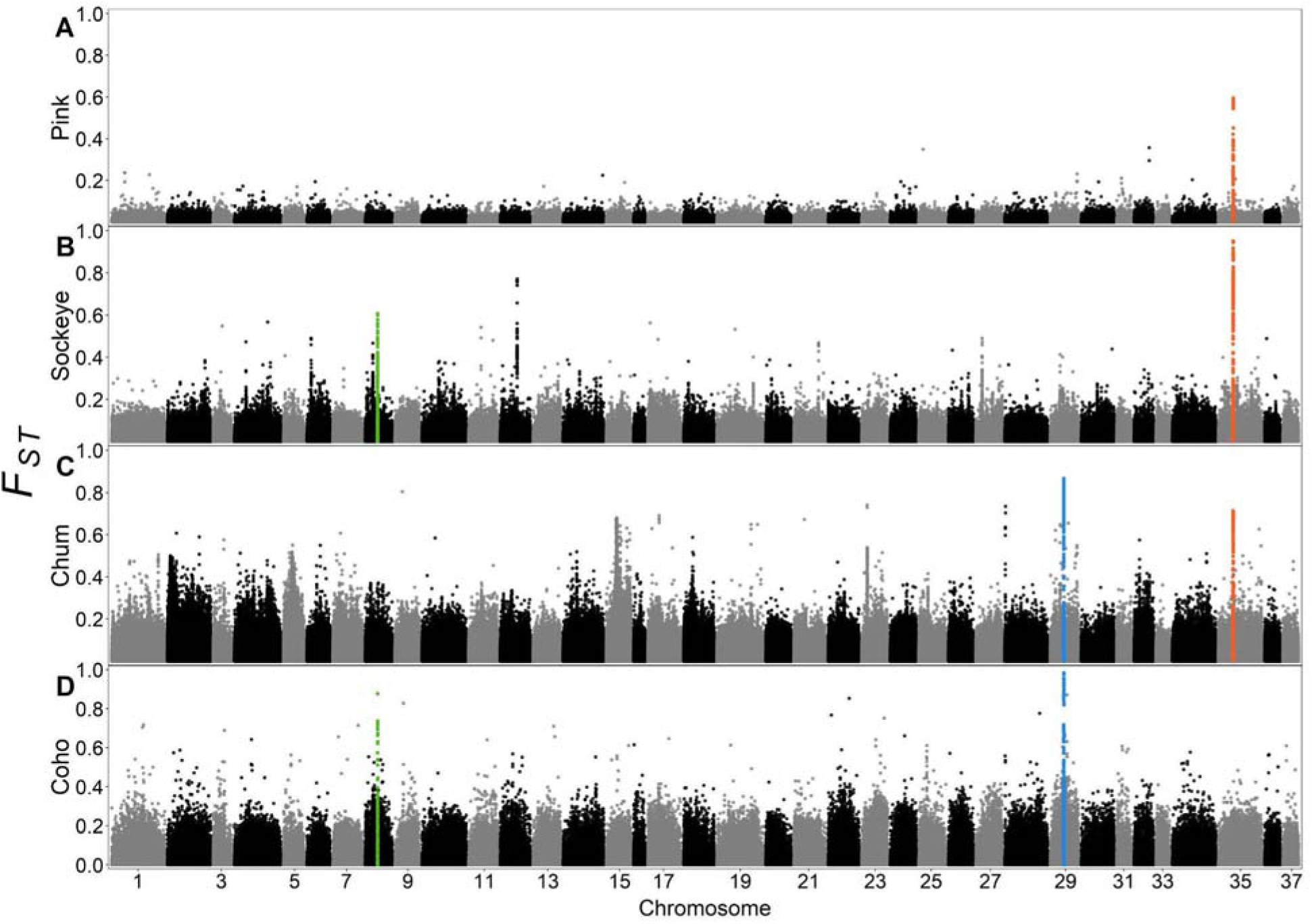
Manhattan plot of genetic divergence (pairwise *F*_ST_) between early and late run fish in four species of salmon. Early- and late-run salmon had high genetic divergence in a duplicated genomic region present on Chromosome 8 (green peak), Chromosome 29 (blue peak) and/or Chromosome 35 (red peak). Pink salmon runs were sampled from the same stream pool and the runs were separated by ∼3 weeks while all other species were sampled in discrete habitats and were separated by 6 weeks to 4 months. All species were aligned to the chum salmon genome. Colors alternate by chromosome.

The three peaks of divergence associated with run timing are ohnologs that arose from the duplication and subsequent diversification of the same genomic region following the 1R/2R and Ss4R WGDs. This region includes numerous transcription factors including several estrogen receptors. Two of these, *esr1* (ERα) and *esr2* (ERβ), originated from a duplication early in the vertebrate lineage (*32*). Within Teleosts, two subtypes of *esr2* have been described, *esr2a* (ERβ*2*, previously ERγ) and *esr2b* (ERβ*1*), which arose during Ts3R (*33*, *34*). The peak on Chr 8 includes the genes *esr1* and spectrin repeat-containing nuclear envelope 1a *(syne1a),* while the peak on Chr 29 spans *esr2b*. The peaks on Chr 29 and Chr 35 occurred within the same small ∼850-kb region of the pre-Ss4R ancestral genome (Fig. 2), which now represents chromosome arms 29a and 35a in chum salmon (homologous chromosome arms referenced in (*16*)). These arms have undergone fusions with other chromosomes and rediploidized since Ss4R and no longer recombine with each other (*30*). The area of highest divergence between early- and late-run salmon on Chr 35 was centered on the leucine-rich repeat containing 9 gene (*lrrc9*), located ∼650 kb away from *esr2b-L*, a duplicated copy of *esr2b* from Ss4R (*-L* denotes *‘like’*). Prior research on sockeye salmon linked variation at *lrrc9* to divergence between stream- and beach-spawning ecotypes (*35*), while *esr2b-L* has been associated with distinct reproductive strategies (single vs. multiple spawning) in steelhead trout (*O. mykiss)* (*36*). Other genes in this region include *six6a*, a gene linked to age-at-maturity and run timing in Atlantic salmon (*Salmo salar)* (*37*, *38*) and age-at-maturity in steelhead (*39*). Additional peaks were present in all species except pink salmon, and were generally unique to each species.

**Figure 2:**
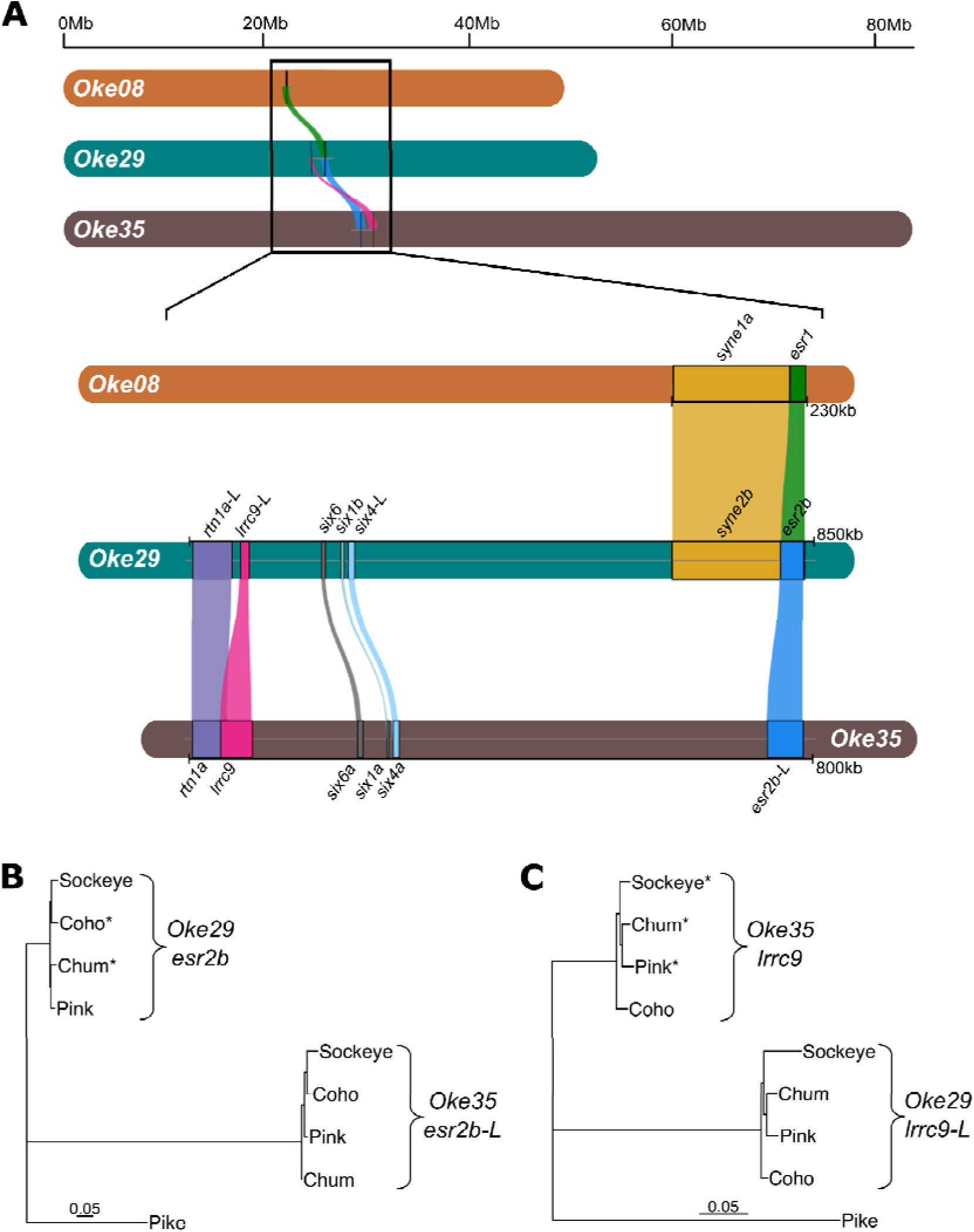
Phylogenetic analysis indicates ohnologs arising from the salmon specific whole genome duplication diverged before speciation. (A) Comparison of an ∼850-kb duplicated region and specific genes associated with run timing on Chromosomes 8 (*esr1*), 29 (*esr2b*) and 35 (*lrrc9*). A previously identified gene associated with age-at-maturity and run timing in Atlantic salmon, and age-at-maturity in steelhead trout (*six6a*) occurs within this region. The orientation of the genes on Chr 35 are opposite of that of Chr 29 because of a large (∼40Mb) and likely ancient inversion that may have played a role in rediploidization. Maximum likelihood trees based on GTR nucleotide substitution model between ohnologs, of (B) *esr2b* and (C) *lrrc9* with Northern Pike (a close relative to the recent common ancestor of salmonids) used as an outgroup. All nodes had 100% bootstrap support. Trees show genes and their duplicated copy (denoted with ‘-L’ for like) grouped by ohnolog rather than by species, consistent with divergence before speciation.

Sequence alignments of the ohnologs on Chr 29 and Chr 35 of pink, sockeye, coho, and chum salmon contextualize the evolutionary history of the ohnologs and provide evidence of species-specific patterns of diversification. The distances between ohnologs were significantly greater than the variation occurring within each gene copy alone (permanova *lrrc9 R^2^* =0.984, *P* = 0.029; *esr2b R^2^* =0.927 *P* = 0.006; Fig. 2B-C) Within gene trees, species grouped together within ohnologs rather than ohnologs of a given species grouping together, indicative of rediploidization prior to species divergence (*20*). For both genes, alleles associated with run timing variation mapped to the same ohnolog. Within each set of ohnologs, sequence divergence from pike was asymmetric, with higher sequence divergence in copies not associated with run timing (*lrrc9-L* and *esr2b-L*) (Wilcoxon test *lrrc9 P* = 0.029; *esr2b P* = 0.029).

There was strong concordance between individual phenotype and genotype with few instances of early homozygous fish with late run timing phenotypes and vice versa (Fig 3, S4). We used PCAs constrained to the highly divergent genomic regions around each shared peak to identify individual genotypes (Fig. 3). Among species for a given gene, early- and late-run genotypes were delineated along the first principal component in the three distinct groups representing homozygous early individuals (possessing two copies of the early allele, EE), heterozygous individuals (one copy of each allele, EL), and homozygous late individuals (two copies of the late allele, LL) (Fig. 3). On average at *lrrc9*, EE individuals were 15 times more likely than LL individuals, and 2 times more likely than EL individuals, to return early (Fig. S4). Similarly, chum salmon with EE genotypes at *esr2b* were 1.3 times more likely to return early compared to EL fish. All run-timing groups had some fraction of individuals that were heterozygous, with two exceptions: the chum salmon late-run collection was entirely homozygous for the late allele of *lrrc9* (Fig. 3C) and the coho salmon early-run collection was entirely homozygous for the early allele of *esr2b* (Fig 3B, S4). Heterozygosity was highest in pink salmon, probably because there is the least temporal separation between run timing groups.

**Figure 3:**
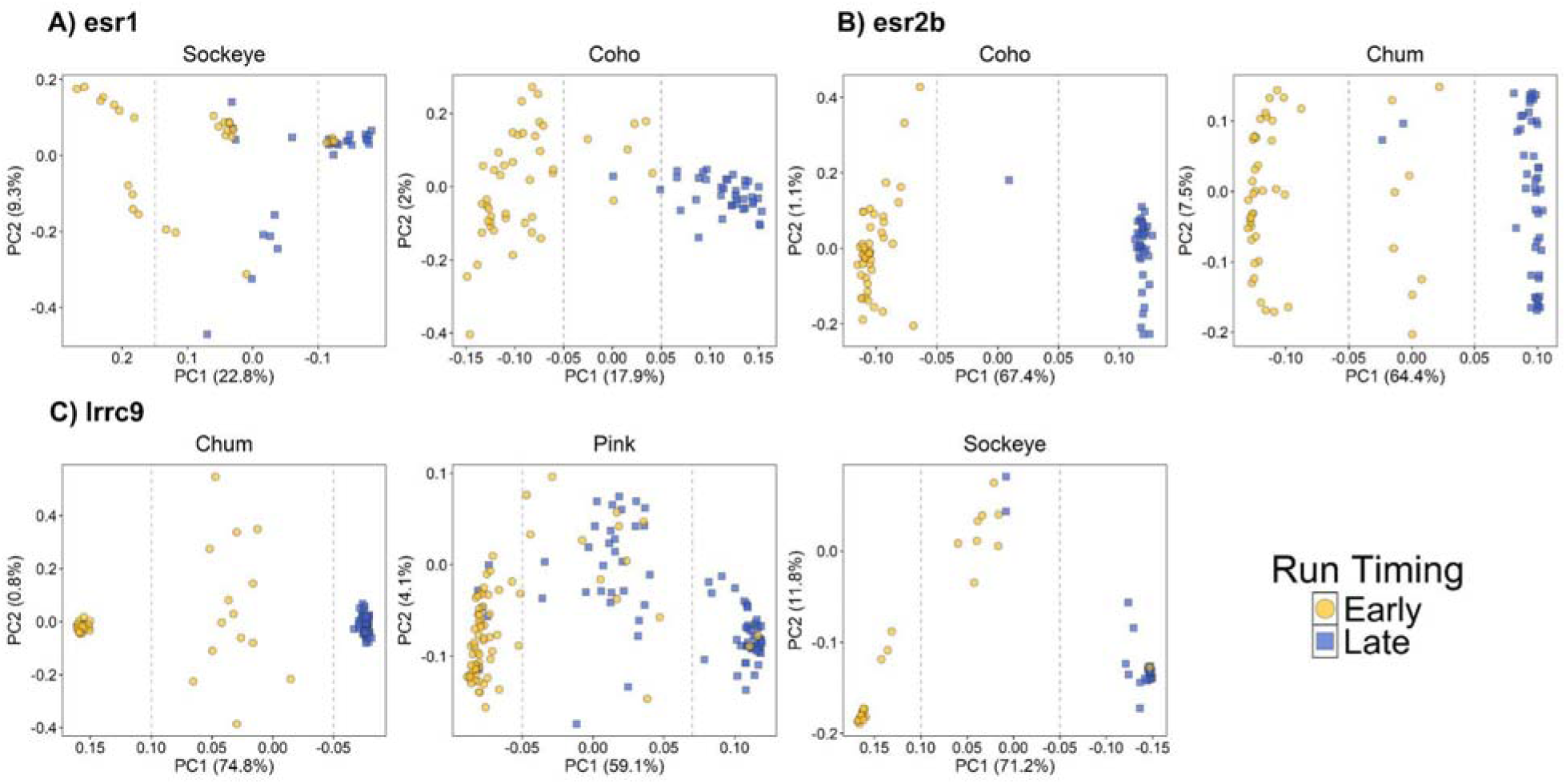
Genotypes at three large effect loci are linked with run timing. Run timing genotypes were identified at three large-effect loci [(A) *esr1*, (B) *esr2b*, and (C) *lrrc9*] with individual based principal component analysis (PCA). PCA of the highly diverged regions for the genes linked with run timing phenotypes and shared across species distinguished early allele homozygotes (left), heterozygotes (center), and late allele homozygotes (right). There was high concordance between run timing phenotypes (color) and genotype (position in PC1 space).

### Alleles evolved independently following species divergence

To determine whether shared trait-genotype associations stemmed from ancestral haplotypes or independent evolution, we compared run timing allele sequences among species. Phylogenetic trees indicated that early and late run-timing alleles at *esr1*, *esr2b,* and *lrrc9* were more similar within than among species, having evolved independently in each lineage (Fig. 4). Divergence peaks were largely distinct among species: *esr1* displayed broad peaks across the 900-kb region in both sockeye and coho salmon; *esr2b* showed a narrow peak within the 1.4-Mb region in chum salmon and a broad peak in coho salmon; and *lrrc9* exhibited the narrowest peak in pink salmon and the widest in sockeye salmon across the 315-kb region (Fig. 5). Across all three gene regions, shared polymorphic sites were exceptionally rare (15 of 816 at *esr1*, 9 of 5317 at *esr2b*, and 21 of 1389 at *lrrc9*), and almost none exhibited substantially elevated divergence (*F*_ST_ > 0.5) in multiple species (Table S3). The near-absence of shared divergent loci, combined with higher divergence among rather than within species, demonstrates repeated allelic divergence after the speciation of the Pacific salmon *∼*6–20 MYA, consistent with other large-effect loci in salmon (*e.g., greb1L*, *six6a;* (*25*, *40*)).

**Figure 4:**
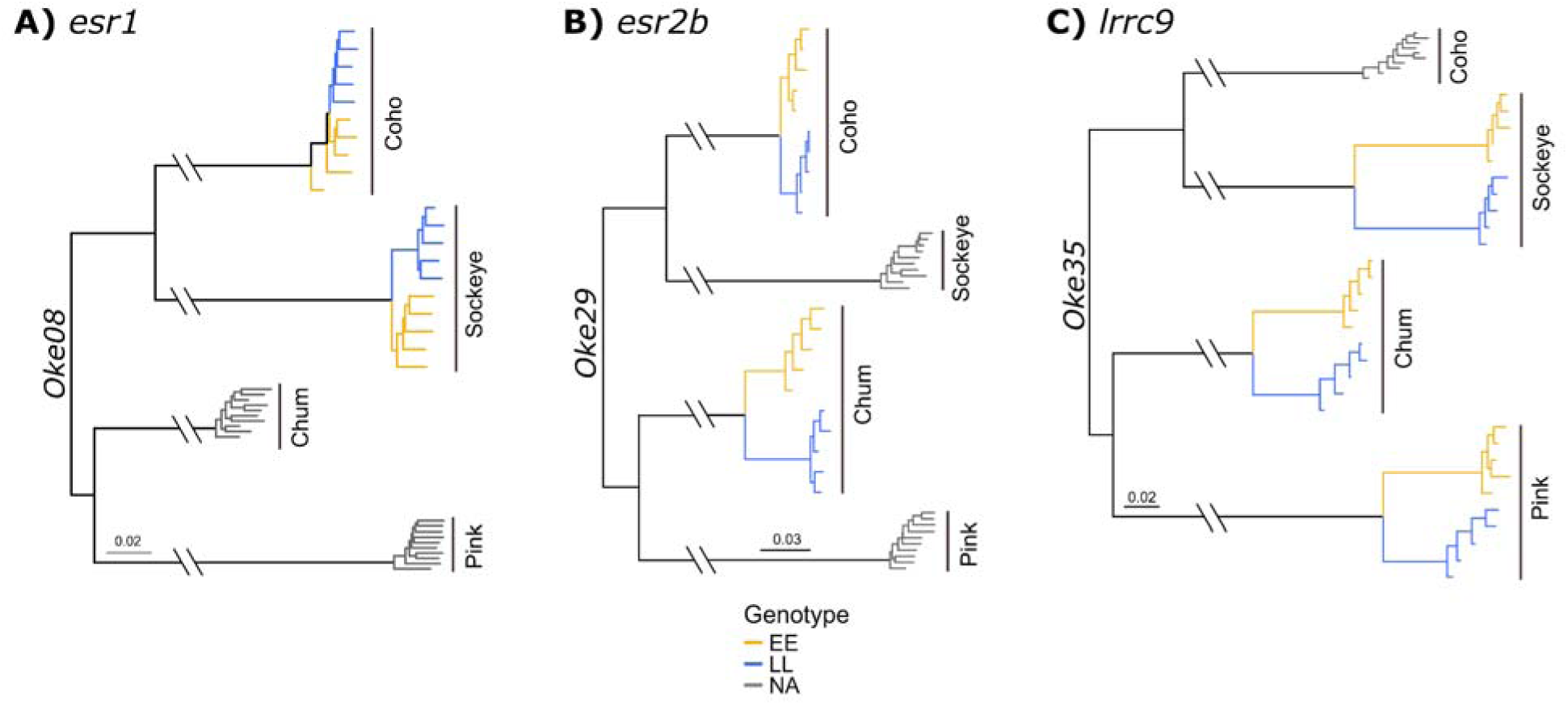
Recurrent evolution of alleles related with run timing at three major effect loci indicates independent evolution of early (EE) and late (LL) alleles in each species following speciation events. Early-and late-run timing alleles at (A) *esr1* on chromosome 8, (B) *esr2b* on chromosome 29 and (C) *lrrc9* on chromosome 35 evolved independently following speciation in Pacific salmon. Phylogenetic trees included ten fish for each species, five individuals from each homozygous genotype defined by PCA (Fig. 3) if associated with run timing and ten individuals with the highest coverage from species that did not show an association. Branch lengths were shortened by 0.130 for *esr1,* 0.091 for *esr2b*, and 0.075 for *lrrc9*.

**Figure 5:**
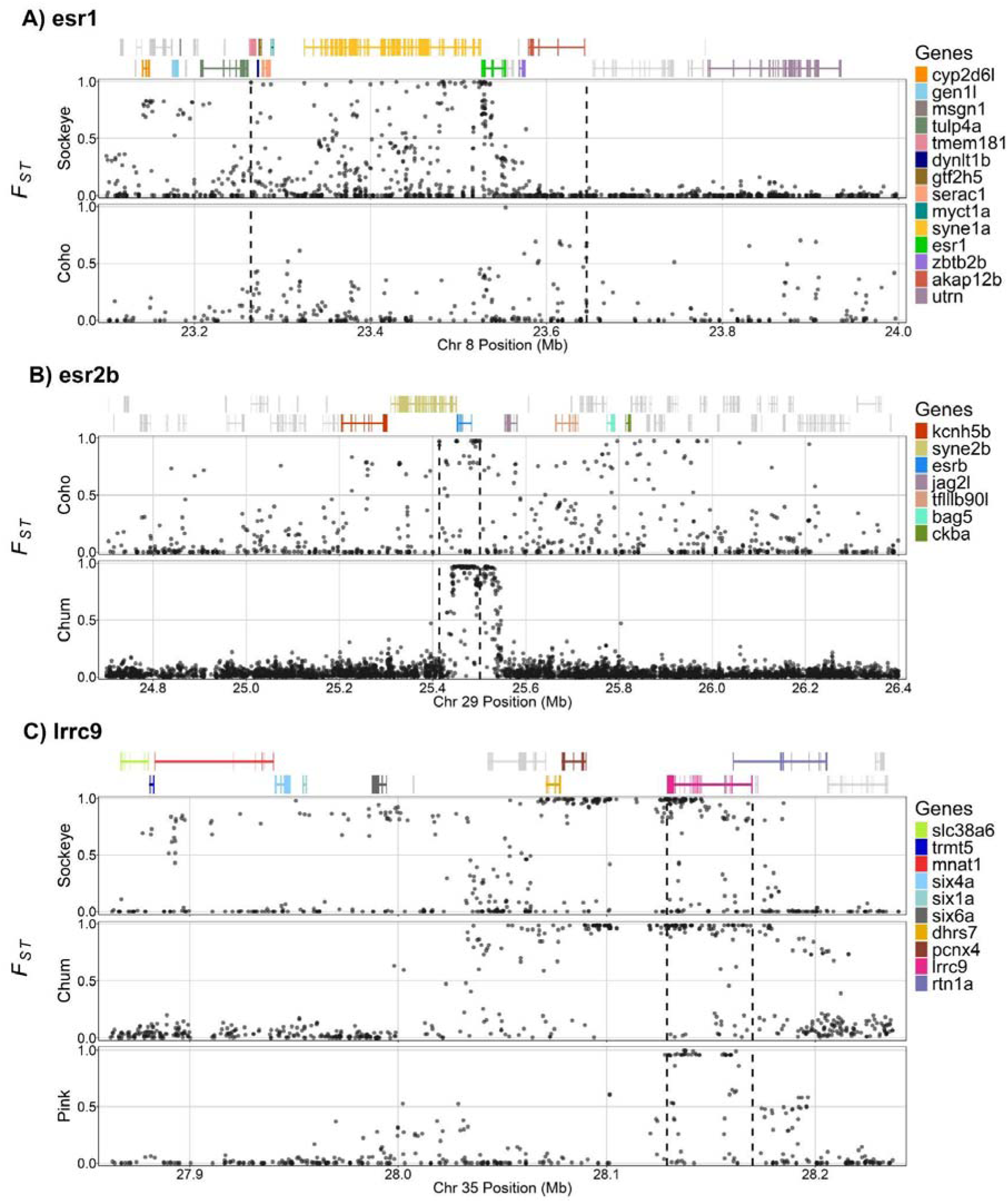
Patterns of genetic divergence between run timing genotypes at three large-effect loci indicate genomic positions most closely associated with run timing. Genetic divergence (*F*_ST_), calculated between EE and LL individuals for (A) *esr1* on Chromosome 8 (B) *esr2b* on Chromosome 29, and (C) *lrrc9* on Chromosome 35. Dashed vertical lines indicate the regions used to construct the PCAs (Fig. 3). Above each plot, genes (lines) and their exons (vertical blocks) from the chum reference genome are shown.

The selective landscape and demographic characteristics within each species likely shaped genetic architecture (*e.g.,* number of loci) underlying run timing variation. Allele frequencies at a single pink salmon locus in *lrrc9* had a strong linear relationship (*r*^2^ = 0.796) with run timing across 21 collections that span temporally isolated lineages (even- and odd-year returns) across much of the species range (Fig. S5). While results from amplicon sequencing targeting variation observed within Auke Creek pink salmon does not preclude the use of additional loci in these collections, it does suggest a conserved mechanism at *lrrc9* for run-timing variation in pink salmon throughout their range. In contrast, sockeye salmon appear to have multiple mechanisms: early- and late-run sockeye salmon that spawned in distinct lake habitats differed at multiple loci, while a single creek differed only at *esr1*. Additionally, Sutherland et al. (*41*) documented an association between run timing and the alternate ohnolog of *lrrc9* (*lrrc9-L*) in the Fraser River. This suggests that sockeye salmon may rely on different run timing loci throughout their distribution, and it demonstrates that each copy retained the latent capacity to generate the major-effect locus at *lrrc9* despite rediploidization prior to species divergence. Pink salmon, characterized by high straying rates relative to other species, may have a simpler genetic architecture because higher rates of gene flow between populations with different selective optima favor few loci of large effect (*42*).

### The duplicated genomic region contains key genes for reproductive regulation

Duplicated genes that undergo subfunctionalization can create multiple distinct ways to fine-tune biochemical pathways (3). We therefore used a KEGG pathway analysis to identify functional systems where genes linked with run timing overlap, highlighting potential pathways under evolutionary modification. Multiple genes within the duplicated region have roles in the hypothalamic-pituitary-gonadal (HPG) axis that are implicated with involvement in estrogen, prolactin and thyroid hormone signaling as well as gonadotropin releasing hormone (GnRH) secretion pathways (Table S8). Given that the HPG axis acts as the master regulator of vertebrate reproductive timing, successive duplication of crucial genes in the HPG axis may have facilitated fine-tuned regulation of development and reproduction.

The duplicated regions linked to run timing contain transcription factors, coactivators, and enzymes that regulate reproduction at different levels of the HPG axis. The ancestral estrogen receptor, having undergone duplication and subfunctionalization numerous times, has resulted in numerous paralogs, each one having distinct functions in the control of reproduction (*43*, *44*). The pathway is driven by the release of GnRH from the hypothalamus, which stimulates the pituitary to secrete hormones that induce gonadal gametogenesis and steroidogenesis. The synchrony in reproductive and migration timing may rely on the detection of photoperiod via the pituitary, which has been shown to modulate the estrogenic regulation of behavior (*45*, *46*). The gene *six6a,* on Chr 35, has been associated with age at maturity and seasonal return timing in Atlantic salmon (37, 38) and age at maturity in steelhead (39). The gene phospholipase C-beta 1 (*plcb1*), on Chr 29, is an enzyme that facilitates many aspects of mouse reproduction including ovulation and embryo development as well as behavior (*47*). The genes *six1* and *six4* are crucial transcription factors involved in gonadal development (*48*). The gene *greb1L*, a paralog and likely ohnolog of the estrogen response gene *greb1* from 1R/2R, is associated with run timing in both Chinook salmon and steelhead (*25*, *40*). The gene *greb1* on Chr 35 and its ohnolog from Ss4R on Chr 29 are ∼11Mb away from the region of highest divergence in this study.

The evolutionary path from multiple WGDs to the diverse migration timing phenotypes that manifest across salmon species is consistent with repeated duplication and diversification. We propose the chronology might have progressed generally as follows: (1) the ancestral estrogen receptor is duplicated during one of the two WGDs early in vertebrate evolution giving rise to *esr1* and *esr2,* (2) these two estrogen receptors subfunctionalize the roles of the ancestral gene, (3) each of these genes are duplicated during Ts3R, resulting in an additional teleost specific estrogen receptor, *esr2a,* and *esr2b,* as well as a duplicated copy of *esr1* that was subsequently lost through nonfunctionalization, (4) the genes *esr2b* and *lrrc9* are duplicated during Ss4R, creating ohnologs on Chr 29 and Chr 35, (5) the chromosomes were functionally tetraploid and recombined, (6) divergence between chromosomes increased and the two chromosomes became functionally diploid, (7) salmon species diverged, and (8) new alleles evolved that contributed to the formation and maintenance of complex migration and reproduction phenotypes in multiple salmonid species.

Although the specific functional differences between the *esr1*, *esr2b*, and *lrrc9* alleles remain unclear, new genetic variation at these loci and their surrounding regions has evolved repeatedly. In salmonids, life-history traits associated with this region have arisen independently at least nine times (*esr1* in sockeye and coho salmon, *esr2b* in chum and coho salmon, *lrrc9* in sockeye, pink, and chum salmon, and *six6a* in Atlantic salmon (*37*, *38*) and steelhead trout (*39*)). Similarly, in Atlantic herring (*Clupea harengus*) and Atlantic cod (*Gadus morhua*) the alternative ohnolog (*esr2a*) arising from the Ts3R duplication has diverged between migration runs (*49*, *50*) (Fig. S6). Remarkably, while this duplicated region makes up ∼0.05% of the salmon genome, it contains roughly half of the large-effect loci or their paralogs. The recurrent association of this region with migratory/maturation phenotypes emphasizes its evolutionary importance.

Given the association with integrated phenotypes that involve multiple physiological and behavioral processes, these large-effect loci may regulate multiple genes. The lack of shared polymorphisms indicates they evolved independently across species to drive new phenotypes. Consistent with other large-effect loci in salmon (*greb1L* and *vgll3*), the strongest associations mapped to intergenic regions. Therefore, these loci may regulate expansive networks (*51*), such as those involved in the HPG axis, or they could harbor an unannotated regulatory element, such as a long noncoding RNA, whose ohnologs span the region (*52*).

By comparing divergence across four salmon species with discrete migration timing, we demonstrated that multiple rounds of WGD and subsequent subfunctionalization of a small set of genes may have been pivotal in the evolution of large-effect loci that underlie variation in reproductive timing. Genes within the region (*esr1* and *plcb1*) have an ancestral role in development across vertebrates. In fish, the Ts3R duplication likely freed one copy of this region to regulate development timing, as seen in cod and herring, while the other copy maintained essential development functions. The subsequent Ss4R duplication in salmonids may allow greater flexibility in timing through greater redundancy of critical development genes, as well as a greater number of targets for mutations. This redundancy may explain why we see many large-effect loci in this region, as other gene copies may buffer deleterious effects of large gene changes. This may have facilitated the successful anadromous life-history of salmonids, where reproductive timing is under particularly intense selection compared to non-migratory fishes and continues to influence their resilience to environmental variation. Future research into the specific patterns of divergence and function of these loci may provide important information on the evolutionary role of WGDs and insights for the management and conservation of salmon.

## Acknowledgments

We thank the numerous graduate students, contractors, and NMFS staff who have provided foundational research and collected samples at the Auke Creek weir, especially John Joyce, Sidney Taylor, Chris Manhard, and Bill Smoker. We also thank the Alaska Department of Fish and Game and U.S. Fish and Wildlife employees for collecting samples and facilitating the transfer of DNA from their archives. We thank Curry Cunningham, Nicolas Lou, Daniel Schindler, Stephanie Carlson, Andrew Piston, and Peter Sudmant for useful discussions about the paper, and Devon Pearse, Robin Waples, and Fred Allendorf for providing valuable comments on manuscript drafts. This work was also supported in part by the high-performance computing and data storage resources SEDNA operated by National Marine Fisheries Service.

## Author contributions

Conceptualization: WAL, DAT, AJG, PDB. Methodology: WAL, DAT, PDB, NH, KDA, GLO, DSB. Investigation: PDB, NH, WAL, DAT, GLO, SV, JR, AB, EL, AJG. Visualization: NH, PDB, WL, DAT. Funding acquisition: WAL, DAT. Project administration: WAL. Supervision: WAL, DAT. Writing – original draft: WAL, PDB, DAT, NH, Writing – review & editing: WAL, DAT, PDB, NH, GLO, DSB, KDA, SV, JR, EL, AB, AJG.

## Data and materials availability

Fastq files for pink, chum, coho, and sockeye (Whitefish and Pick creeks) are available through NCBIs Sequence Read Archive (SRA; BioProject: PRJNA1259009; SRA submission: SUB15296226). Sequences from sockeye populations Teal Creek and Anvil Beach are available at NCBI SRA (PRJNA1006708). All code and non-sequencing data are available at https://github.com/AFSC-Genetics/Salmon_Run_Timing.

## Materials and Methods

### Study locations and sample collections

We obtained genetic samples from pink (*n_even_* = 153, *n_odd_* =135; see below for description of even and odd lineages), sockeye (*n* = 196), chum (*n* = 96), and coho (*n* = 96) salmon displaying variation in run timing (Table S1, Fig. S1). All individuals were of unknown age, except for pink salmon, which are obligate 2-year-olds. Each species was collected in a different region of Alaska, which included Southeast Alaska near Juneau, lower and upper Yukon River in northern Alaska and Canada, Bristol Bay, and the Kenai Peninsula (see Fig. S1).

Pink salmon in our study were collected from Auke Creek, a small creek near Juneau in Southeast Alaska. This stream supports early and late runs of pink salmon, the modes of which are separated by approximately 20 days (*53*). The stream has been extensively studied, and differences in run timing and developmental rates of early- and late-run pink salmon in Auke Creek are partially due to additive genetic variance (*54*, *55*). Pink salmon have an obligate two-year life cycle, resulting in genetically distinct odd- and even-year lineages that spawn in the same reaches in alternating years at Auke Creek. We obtained samples, axillary processes stored in ethanol, of early- and late-run pink salmon, defined by their return timing through a permanent weir, from the odd- and even-year lineages from a previous study (*56*). We selected 288 of these samples for sequencing, with equal representation among run timing, lineages, and sexes (Table S1). Compared to the entirety of the return, the odd-lineage early samples were collected at an intermediate time and therefore did not represent the earliest returning spawners in those years. The resulting whole genome scan did not exhibit a peak of divergence as large as that of the even-year lineage (see Fig. S2A). To investigate patterns in the odd-year lineage further, additional samples were obtained from the 2019 return to the Auke Creek Weir (Table S1) and analyzed for variation at *lrrc9* (see *<u>Design of a high throughput assay for pink salmon</u>* section below).

Sockeye salmon in our study were collected from the Wood River drainage, a well characterized watershed inhabited by sockeye salmon on the west side of Bristol Bay, where differences in migration timing of the distinct populations can span 2.5 months and are associated with spawning habitat (*28*). We collected samples, fin clips stored on Whatman paper, from one lake beach and three streams in the Wood River drainage that are approximately 20–40 km apart: (i) Teal Creek, an early-spawning stream that peaks in early to mid-August, (ii) Whitefish Creek, a late-spawning stream that peaks near the end of August, (iii) Anvil Beach, a late-spawning beach that peaks around mid-September, and (iv) Pick Creek, an early spawning stream that peaks in early to mid-August with two separate collections, one at the beginning and one at the end of the run (*57*). Whole genome sequencing data from Anvil Beach and Teal Creek were originally used in Euclide et al. (*58*) with an equal sex ratio whereas Whitefish Creek samples were all males (collected for an ongoing study comparing males of different ages). Comparisons of divergence among sockeye populations are between Teal Creek (early) and Anvil Beach (late) unless otherwise noted.

Chum salmon in our study were collected from the lower Yukon River in Alaska and the upper Yukon River in Canada. The Yukon River has summer spawners, which enter the river in early summer (June - July) and spawn soon after in the lower reaches, and fall spawners, which enter the river in mid-summer (July-August) but swim as far as 3190 km from the mouth of the river to spawn in the upper reaches in fall (*59*). Fall spawners are generally larger, have larger lipid reserves, and enter the river less mature than summer spawners (*59*). We obtained extracted DNA from Yukon River summer- (Gisasa River) and fall- (Pelly River) run chum salmon, from individuals sampled in spawning stretches of each river. These populations spawn approximately two months apart and are separated by more than 1500 river km.

Coho salmon in our study were collected from the Kenai River drainage, which is located in Southcentral Alaska near the northern end of the coho salmon range and has an uncommonly protracted run, especially for a salmon population at such high latitude (*26*). We obtained extracted DNA from fish sampled in spawning habitat in both a tributary of Skilak Lake (Skilak River) and the Kenai River mainstem just below the outlet of Skilak Lake. These populations are separated by about 30 km and spawn approximately four months apart.

Although these collections capture diverse run timings across the four species, further sampling across their geographic ranges is needed to determine how broadly these associations are conserved.

### Generating Whole Genome Data and Initial Filtering

We conducted whole genome resequencing on all samples except 54 sockeye, which were previously sequenced in Euclide et al. (*58*). Tissue samples (fin or axillary process clips) were either dried or preserved in ethanol and DNA was extracted with Qiagen DNeasy Blood and Tissue Extraction kits (Hilden, Germany) or Machery-Nagel NucleoSpin tissue core kits (Duren, Germany) at the NOAA Laboratory in Juneau, Alaska or the Alaska Department of Fish and Game Laboratory in Anchorage, Alaska. All sequencing libraries were prepared following Therkildsen and Palumbi (*60*), and all libraries were sequenced with paired-end 150-bp (300 cycle) chemistry on Illumina sequencers. DNA concentration was normalized to 10ng based on PicoGreen fluorescence measured on a BioTek Synergy HTX microplate reader and DNA quality was visualized on a 2% agarose gel. Tagmentation was performed with Nextera DNA Library Prep Kit (Illumina, Inc., San Diego, CA). Illumina adapter sequences and sample barcodes were added to tagmented DNA fragments by low cycle PCR with KAPA HiFi HotStart polymerase. PCR products were then normalized to 25ng with SequalPrep plates (ThermoFisher Scientific, Waltham, MA, USA) and pooled into a single library. The normalized pooled library was purified and size-selected with 0.6x AMPure XP beads. Libraries were then visualized on an E-gel and quantified using a Qubit HS dsDNA Assay Kit (ThermoFisher Scientific).

Chum and coho salmon samples were sequenced with Illumina Novaseq X 10B chemistry on a single lane for each species (96 samples per lane) at Seqmatic (Fremont, CA). Sockeye salmon samples from Whitefish Creek were sequenced on a single Novaseq 6000 S4 lane at the University of Oregon Genomics and Cell Characterization Core Facility. Samples from Teal Creek and Anvil Beach (available from (*58*)) were also sequenced on a Novaseq 6000 S4, but at higher coverage (32 individuals per lane). Samples from Pick Creek were sequenced on a single Novaseq X Plus 25B lane at SeqMatic. Pink salmon samples were sequenced on a Novaseq 6000 S4 (96 individuals per lane) at Novogene (Sacramento, CA).

Sequencing files were trimmed of adapters with TRIMMOMATIC v0.39 (*61*) and poly-G tails with the program fastp using the cut_right flag (*62*). Trimmed sequences were aligned with BWA v0.7.17 to the chum salmon reference genome (GCF_023373465.1_Oket_V2_genomic (*25*)), a phylogenetically intermediate species (*63*), in order to directly compare genomic regions of divergence across species. SAMtools (*64*) was used to convert between file formats. PCR duplicates were removed with picard v2.23.9 (http://broadinstitute.github.io/picard/) and BamUtil v1.0.5 (*65*) was used to clip overlapping read pairs. Sequencing depth for each individual was calculated with SAMtools v1.11 (*64*) and samples with mean sequencing coverage that fell below 0.4x were removed from analyses. Within each species, average sequencing depth was compared using a Welch’s *t*-test (□= 0.05) across each run timing phenotype. Only the even-lineage pink salmon had statistically different average depths, and a subset of BAM files from the higher-coverage group were subsequently downsampled with SAMtools (*64*) to match the lower-coverage group’s average depth. Although the sockeye salmon samples from Euclide et al. (*58*) had deeper sequencing coverage than samples from Whitefish Creek, the effective sample size (*66*) was approximately equivalent because Whitefish Creek had more samples (*n* = 48 vs. *n* = 27) but lower average depths (1.6x vs 4.7x); therefore, sockeye salmon samples were not downsampled. Additional information on sample sizes and sequencing coverage is available in Table S1.

### Genetic Divergence Associated with Run Timing

Single nucleotide polymorphisms (SNPs) were identified and genotype likelihoods (GLs) were estimated for each species with ANGSD v0.933 (*67*) using the SAMtools genotype likelihood calling algorithm. Each species had the following filters: GL = 1, doMaf = 1, uniqueOnly = 1, remove_bads = 1, only_proper_pairs = 1, C = 50, minQ = 20, minMaf = 0.05, snp_pval = 1e-10, and minInd = 0.3*sample size (N). Since all species were aligned to the chum salmon reference genome, minMapQ was set to 20 for chum samples but lowered to 15 for all other species to account for increased mapping errors associated with alignment to a different species. For species with average depth exceeding one, minDepth = N and maxDepth = 20*N; otherwise, minDepth = ½*N and maxDepth = 10*N. The resulting number of SNPs across the genome after filtering averaged 3.1 million (1.1 million for coho to 6.3 million for chum). Additionally, species-specific reference genome alignments for pink (GCF_021184085.1_OgorEven_v1.0_genomic), sockeye (only Teal and Anvil samples; GCF_006149115.2_Oner_1.1_genomic), and coho (GCF_002021735.2_Okis_V2_genomic) were used to evaluate reference genome bias (Fig. S7). The alignments follow the above methodology and genotype likelihood filtering except that minMapQ was increased to 20 for all species and the minInd flag was not included for coho.

Divergence at each polymorphic site between early and late run timing phenotypes was calculated with *F*_ST_ (fixation index), a metric that ranges from 0 to 1, where values of one represent complete divergence. For pairwise *F*_ST_ scans, the minQ, minMaf, and snp_pval flags were removed because the polymorphic sites file was included with the -sites flag. We used the folded site frequency spectrum with ANGSD’s realSFS because the ancestral state was unknown (early or late). Manhattan plots of *F*_ST_ across the genome, aligned to the chum salmon reference, were plotted for each species with R statistical software (v.4.3.2; R Core Team 2021) and visualized with ggplot2 (*68*). Similarly, Manhattan plots of *F*_ST_ across the genome were plotted for species-specific alignments. Global and genomic region weighted *F*_ST_s were calculated as the ‘ratio of averages’ of polymorphic sites as defined in Table S2 (*69*). A local score approach was used to determine the significance of regions with elevated *F*_ST_ (*31*). Allele count data from early and late run timing collections of each species, aligned to the chum reference genome, were compared with a Fisher’s exact test from which significance values were used as input for the local score analyses (ξ = 3, □= 0.01). The selection of a tuning parameter (ξ), for which we chose a conservative value, is a tradeoff in power to detect true outliers associated with run timing and background variation, or variation associated with spawning location, particularly in the cases of chum, coho, and sockeye salmon collections. Shared peaks among species, defined as those within 100 kb of the local score boundaries, were identified (Table S4).

After regions of elevated *F*_ST_ were identified with local score, species-specific alignments were used to identify SNP positions and determine amino acid substitutions. SNPs from seven regions for the four species were evaluated: pink salmon genes within the *lrrc9* region on chromosome 10 (NC_060182.1:72650536-72875058), chum salmon genes within the *esr2b* region on chromosome 29 (NC_068449.1: 25440358-25545548) and the *lrrc9* region on chromosome 35 (NC_068455.1:28033205-28206418), sockeye salmon genes within the *lrrc9* region on chromosome 12 (NC_042546.1:41162738-41448689) and *esr1* region on chromosome 13 (NC_042547.1:7206200-7643690), and for coho genes within the *esr2b* region on chromosome 7 (NC_034180.2:29455765-30509378) and *esr1* region on chromosome 21 (NC_034194.2:24716700-24902281). Genes within these regions were extracted from the genome general feature format (.gff) file of each species, and their sequences were extracted from the genome with the R package and function biostrings::readDNAStringSet (v2.78.0). For each transcript variant of the gene within a species, polymorphic sites identified with ANGSD and with *F*_ST_ exceeding two standard deviations above the mean for a given chromosome were transcribed with the R package and function seqinr::translate (v1.0-2) and evaluated to determine if the SNP changed the transcribed amino acid. For the 72 genes evaluated, there were 1,962,014 sites of which 1686 were polymorphic. Averaged over the transcript variants of a gene within a species, 96.3 sites were polymorphic in the coding DNA sequence (CDS) (5.7%) and 31.8 (1.8%) caused non-synonomous mutations.

Convergence of early and late protein variants among species was evaluated for all genes in which there were non-synonymous mutations in multiple species: *lrrc9*, *rtn1a*, and *syne2b*. At *lrrc9*, pink salmon had two non-synonymous mutations: a change from leucine (L) in late run fish to valine (V) in early fish at codon 280 and from glycine (G) in early fish to serine (S) in late fish at codon 1392. While an L to V mutation is a conservative change of one nonpolar hydrophobic amino acid for another, the change of G to S substitutes a small, nonpolar and flexible amino acid with a larger, polar amino acid. This non-conservative change may increase polarity and potential phosphorylation sites. Sockeye salmon had a single non-synonymous mutation: a change from methionine (M) in late run fish to isoleucine (I) in early fish at codon 1473, a conservative substitution that replaces one hydrophobic amino acid with another. At *rtn1a*, in chum and pink salmon there is an aspartic acid (D) to tyrosine (Y) and glutamic acid (E) respectively. While the D to E change is conservative, the D to Y substitution is non-conservative, changing a small negatively charged hydrophilic residue with a large polar, hydrophobic one. At *syne2b,* there was a single nonsynonymous mutation in chum salmon leading to a nonconservative change from lysine (K) to asparagine (N) for multiple transcription variants of the gene. While both are polar, K is positively charged and N is uncharged. Coho salmon displayed many more nonsynonymous mutations, up to five depending on the isoform, between run timing groups (Table S5). Three of these were nonconservative, two were conservative, and one in two isoforms terminated the protein a single amino acid early. None of the nonsynonymous mutations were shared among species and there was larger divergence between than within a species. For instance, at *lrrc9* there were 11 amino acid differences between early migrating pink and sockeye salmon.

### Investigating the duplicated regions of association and constructing gene trees of *esr2b*, *lrrc9*, and their duplicated variants

Genome scans revealed three shared regions of elevated *F*_ST_ associated with run timing on chromosomes 8, 29, and 35. Genes within this region have been duplicated and retained after multiple rounds of whole genome duplication. From the salmon specific WGD, arms on chromosomes 29 and 35 are duplicated versions of the same ancestral chromosome and include diverged copies of the same genes (*30*). These gene regions have diverged sufficiently to allow short-read sequences to map to their respective chromosomes. To further ensure paralogous loci were accurately separated and to prevent reads from unresolved duplicates from stacking on a single gene copy, we filtered for high mapping quality and applied a strict maximum read depth filter of 20 times the number of individuals, excluding any loci with abnormally high coverage. A small portion of this ancestral chromosome had an estrogen receptor and synaptic nuclear envelope genes that were duplicated early in the vertebrate lineage. To identify patterns of homology between chromosomes 29 and 35, we used blastn to compare the chromosomes with default parameters (*70*). This revealed a large inversion on chromosome 35 relative to chromosome 29. We visualized the ∼850-kb duplicated region on chromosomes 29 and 35 in the R package geneviewer v0.1.10 (*71*).

We also constructed phylogenetic trees of all the ohnologs with one copy linked to run timing to investigate their evolutionary history. Protein sequences for all estrogen receptors from the four species as well as representative species that were subject to Ts3R but not Ss4R duplications, as well as non-teleosts which were retrieved from NCBI (Table S6). Protein sequences were aligned with Clustal Omega v1.2.2 (*72*) with three refinement iterations, and evaluating the full distance matrix for initial and refinement guide trees. A maximum likelihood tree was constructed with the R package ape v5.8 with the JTT amino acid model and node support from 100 bootstrap replicates (*73*) (Fig S7). For *lrrc9, esr2b,* and their ohnologs *lrrc9-L and esr2b-L (*genes predicted *in silico* during genome annotation with a high degree of similarity to known genes are assigned the same name with a ‘*L*’ for ‘like’), gene sequences for each species were downloaded from its reference genome on NCBI (Table S7). Sequences were aligned with Clustal Omega v1.2.2 (*72*) with default parameters. Alignments of *lrrc9* had a mean length of 24,546 bp with 41.2% pairwise identity and 17.2% identical sites. Alignments of *esr2b* had a mean length of 34,770 bp with 28.4% pairwise identity and 5.4% identical sites. A maximum likelihood tree was constructed with the R package ape v5.8 and the GTR nucleotide model (*74*) with 100 bootstrap iterations and 50% threshold support. Both trees were imported into R with the ape package v5.8 (*73*) and visualized with ggtree v3.10.1 (*75*). Within each gene tree, pairwise distances were calculated with ape v.5.8 (*73*). Sequence divergence between ohnologs were compared with a permanova test in vegan v.2.7 (*76*) and heterogeneity in mutational rate from the outgroup pike was accessed with a Wilcoxon test.

### Characterizing region associated with run timing on chromosomes 8, 29 and 35

To define early- and late-run timing alleles and characterize patterns of divergence within and among species within the duplicated region associated with run timing, we defined large-effect locus boundaries and inferred individual genotypes. Boundaries around the large-effect loci on which to conduct PCA were determined by identifying the highly diverged regions linked with run timing phenotypes (see above) that were shared across all species. Regions that were only divergent in a comparison of a single species were omitted. For chromosome 8, we selected a broader boundary that exceeded the *esr1* gene boundary (NC_068428.1:23264160-23645630), for chromosome 29, we selected a boundary that slightly exceeded the *esr2b* gene (NC_068449.1:25414060-25501622), and for chromosome 35, we selected the *lrrc9* gene as the boundary (NC_068455.1:28128954-28169980). These boundaries were used consistently across all species. PCAs were constructed using GLs within the boundaries to estimate covariance matrices between individuals with PCAngsd (*77*). Individuals were assigned a genotype based on which of the three distinct clusters across PC1 they grouped into: homozygous early (EE), heterozygous (EL), and homozygous late (LL).

To better evaluate the SNP sites in *esr1, esr2b*, and *lrrc9* that were most divergent between run-timing genotypes, while accounting for genotype by phenotype mismatch, we conducted localized *F*_ST_ scans between homozygous genotypes (EE vs. LL) for each species. We created Manhattan plots, including sites up to 1 Mb away from the gene boundaries to ensure that the full extent of the regions of association were identified (Fig. 5). For sockeye, we incorporated homozygous individuals from the late returning stream population, Whitefish, with the homozygotes from Teal Creek and Anvil Beach to minimize the influence of spawning habitat. After examining divergence at each variant, we defined the regions of association as the genomic span with an *F*_ST_ > 0.5 between run timing alleles in at least one species (NC_068428.1:23100059-23997201 for *esr1*, NC_068449.1:24838997-26208328 for *esr2b*, and NC_068455.1:27877276-28192281 for *lrrc9)*. The regions were defined based on the broadest peak across any species (*e.g.*, sockeye had the broadest peak at *lrrc9* therefore all *lrrc9* visualizations across species use the sockeye boundary). Genes and corresponding exons within the region were positioned using the chum reference genomic feature format file (GCF_023373465.1_Oket_V2_genomic.gff), and characterized genes that contained at least one SNP with *F*_ST_ > 0.5 were labeled in the legend. We identified gene ontology terms for all genes characterized in the corresponding regions using the gene association file of the reference genomes and derived corresponding gene functions from the available coho salmon gene annotations database using Ensembl’s biomaRt (Table S3 (*78*)).

In order to determine if genes within the shared regions affect shared biological pathways, a Kyoto Encyclopedia of Genes and Genomes (KEGG) analysis was performed. The R package KEGGREST (*79*) was used to recover the KEGG Orthology (KO) identifiers of each gene and reconstruct the pathways. Most genes within the regions were present in the KEGG database, however *lrrc9* was not. The most frequent class of network was organismal systems; endocrine systems, with estrogen, prolactin, and thyroid hormone signaling as well as GnRH secretion pathways that have multiple genes within the regions present (Table S8). These pathways play important roles in the hypothalamic-pituitary-gonadal (HPG) axis which is fundamental for regulating reproduction and development and often have the genes *esr1*, *esr2b* and *plcb1* present. Also, the genes *mnat1* and *gtf2h5* were present in pathways associated with transcription and nucleotide excision repair. As part of transcription factor IIH, these genes can act as a coactivator of estrogen receptors by facilitating their phosphorylation (*80*). While not included in the KEGG analysis, genes on chromosome 28 (*ngfa* and *thyrotropin*) that had diverged between early and late returns but failed to meet criteria for inclusion for both chum and coho salmon (significant in chum but not in coho), function at multiple levels of the HPG axis. Nerve growth factor can stimulate GnRH neuron differentiation and release in the hypothalamus, enhances gonadotropin synthesis and secretion in the pituitary, and influences ovulation and spermatogenesis in the gonads (*81*).

### Evolutionary origins and divergence of run-timing loci across species

While run timing is associated with the same major-effect loci across multiple species, the size, divergence, and specific polymorphic sites of these regions of covariation differ considerably among them. Across the 897-kb region near *esr1*, there were 816 polymorphic sites across sockeye and coho salmon, 15 of which were polymorphic in both species (Table S3). The peak in sockeye, as compared to coho, was characterized by higher *F*_ST_ values over a broader region and encompassed more genes (Fig. 5A). This difference was also present at the 1.4 Mb region near *esr2b,* where nine of the 5317 polymorphic sites were shared between comparisons of early and late alleles of chum and coho. Chum salmon displayed a narrow peak centered around *esr2b* and *syne2b* as compared to the broad peak observed for coho salmon (Fig. 5B). Across the 310-kb region near *lrrc9,* there were 1389 polymorphic sites, of which 21 were polymorphic in multiple species. Pink salmon had the narrowest peak while sockeye had the widest (Fig. 5C). Compared to pink and chum salmon, sockeye salmon displayed high divergence (*F*_ST_ > 0.5) downstream of *lrrc9,* located to the left as *lrrc9* is transcribed on the complement in the chum reference, within an additional five genes (*six6a, six1a, mnat1, trmt5,* and *slc38a6;* Table S2). Polymorphic sites were overwhelmingly unique to a single species; only two shared SNPs at *lrrc9* had *F*_ST_ > 0.5 between early and late alleles of sockeye and chum (Table S2).

Phylogenetic trees of run timing alleles were used to explore sequence homology and divergence within and among species. To describe the relationships among the genomic regions most linked to run-timing variation, we constructed trees based on the shared highly divergent regions; NC_068428.1:23100059-23997201 (7211 SNPs) for *esr1*, NC_068449.1:25414060-25501622 (1899 SNPs) for *esr2b*, and NC_068455.1:28128954-28169980 (888 SNPs) for *lrrc9*. These smaller regions were used to mitigate the influence of species-specific markers in regions with lower association that are not necessarily associated with run timing. We calculated pairwise identical-by-state distances with ANGSD by sampling a single read from each site for ten samples per species (-doIBS 1 -makeMatrix 1). We subset the individuals to ten per species, five per homozygous group (EE and LL), selecting individuals with the highest average depth of coverage in each group. For species that did not display divergence at a particular gene region (*e.g.*, *esr2b* in pink, *lrrc9* in coho), we included the ten individuals with the highest depth of coverage. We created neighbor-joining trees with ggtree (*75*). If alleles arose after speciation, early and late homozygotes of the same species should be more similar to each other than to early and late homozygotes of other species. Alternatively, if alleles arose before speciation, early and late alleles should be more similar among species.

### Investigating the relationship between run timing alleles and run timing phenotypes

We investigated the relationship between run-timing phenotype and genotype. Phenotype (early or late) was defined by date of return of the sample collection relative to other collections of the same species. Genotype for each major-effect locus was inferred from local PCA (see above) and from a high-throughput assay for odd-lineage pink salmon (see below). Associations between genotype and phenotype were made by plotting PCAs by phenotype (Figs. 3 and S3) and barplots of the proportion of individuals assigned to each genotype (Fig. S4). Effect size of each genotype was calculated as a relative risk with outcome as returning early or late and exposure being the genotype of the individual (EE vs. LL and EE vs. EL). Due to the sampling design, number of individuals, and depth of sequencing coverage, an effect size from genome-wide association with run timing could not be made across all species.

All species had high concordance between genotype and run timing phenotypes (Figs. 3 and S3). At *lrrc9* for pink, chum, and sockeye salmon, the vast majority of individuals with early returning phenotypes had EE genotypes (left side of PCA; Fig. 3C): 57% of chum, 81% of pink salmon even-lineage, 67% of sockeye salmon (Figs. 3C and S4). All early collections had some proportion of heterozygote genotypes: 30% chum, 16% pink (even-lineage), and 30% sockeye salmon (Fig. S4). While all early collections had some low frequency of late allele homozygotes, there were no early allele homozygotes in either the chum or sockeye late beach collections and only two heterozygotes (7%) observed in the late beach sockeye collection. The late run chum salmon population was fixed for the late run timing allele.

Genotypes at *esr1* for both sockeye and coho salmon were linked with phenotype (Figs. 3A and S3A). The early coho salmon collection was mostly early allele homozygotes (87%) and the late collection was nearly all late allele homozygotes (95%). There were no homozygotes that had the alternate phenotype (Fig. S4). In sockeye, the early Pick Creek collection was mostly EE genotypes (77%), the late Pick Creek collection had only two EE genotypes (4%), and the late beach samples were mostly LL genotypes (63%). However, the association was weaker in Teal Creek (early stream) and Whitefish Creek (late stream). The early Teal Creek sockeye collection had the lowest proportion of EE genotypes (41%) of all early collections, and no LL genotypes were observed in the late Whitefish Creek collection.

Chum and coho salmon had, on average, a stronger association between genotype and phenotype at *esr2b* than either *lrrc9* or *esr1*. Coho salmon early and late collections were nearly fixed for their respective *esr2b* run timing alleles and only a single heterozygous coho salmon was observed in the late run (Fig. 3B). The early- and late-chum salmon runs had low frequencies of heterozygotes, 17% and 4% respectively. In general, more heterozygotes were observed in early run collections, but pink salmon were the major exception, and fewer heterozygotes were observed at *esr2b* than at *lrrc9* and *esr1* (Fig. S4). The populations of coho and chum salmon in this study have greater temporal separation in run timing and larger geographic distance between spawning grounds than pink and sockeye salmon populations (see Fig. S1), which may influence the lower frequency of heterozygotes. The lack of heterozygotes in these collections could also reflect sampling at the extremes of their return (e.g., the earliest returning chum salmon of summer run Gisasa River and the latest returning fish to fall run Pelly River).

Weak genetic divergence between categorical run time collections in pink (odd-lineage) and sockeye salmon (early vs. late stream) demonstrated that return timing of populations can be continuously distributed and likely varies in major-effect locus allele frequency throughout the return. In both comparisons, genetic divergence was low between run timing groups (Fig. S2). Local PCAs at *lrrc9* (from whole genome data) for odd-lineage pink salmon had only early run timing phenotypes present in the putative early homozygous cluster, but early individuals were also found in the heterozygous and late homozygous clusters (Fig. S3C). Sockeye from the late stream had both homozygous and heterozygous genotypes in similar proportions (Figs. S3B and S4). In both cases, the collection of samples representing the categorical (early vs. return) return timing may not be representative. The high prevalence of early pink salmon in the late clusters was likely a result of sampling from the middle and end of the run without sampling the earliest returns. Similarly, the late stream sockeye population from Whitefish Creek were not collected at the end of the run. The high proportion of heterozygosity observed is possibly due to collecting samples during the middle of the run, although this has yet to be confirmed with additional sampling.

### Design of a high throughput assay for pink salmon

To investigate the weak divergence between early and late runs of pink salmon in the odd-lineage, and address possible sampling of intermediate run timing classified as the ‘early’ component, a high throughput assay was developed to genotype samples for *lrrc9*. Pink salmon samples from both even- and odd-lineages were aligned to the pink salmon genome (GCF_021184085.1) and analyzed for divergence between run timing with the same filtering and analysis methods applied to all species aligned to the chum reference genome. Consistent with results from alignments to the chum salmon reference genome, the *lrrc9* peak region was more prominent in the even-lineage, yet a small peak was also present in the odd-lineage. The region of the genome within *lrrc9* with highest *F*_ST_ between early and late groups was targeted for development of a high throughput genotyping panel. PCA was used to identify early and late *lrrc9* alleles and SNPs within the region of highest divergence were targeted for development. After identifying SNPs, 150 bp before and after the SNP were extracted from the reference genome. Primers were developed with Primer3 (*82*) implemented in Geneious Prime 2023.2.1. For each SNP sequence, a 130–170 bp region was targeted with a product size of 100–120 bp. Selected primers had the following characteristics: size 18–25bp, T_m_ 57–63 (max difference 6), and %GC 20–80. Sequences for which primers could not be designed were discarded. Of the successfully designed primers, we developed an assay for the locus OgoE1_LG10_72721832, named after the reference genome used (OgoE1; GCF_021184085.1), the linkage group (Chromosome 10) and position (72721832):

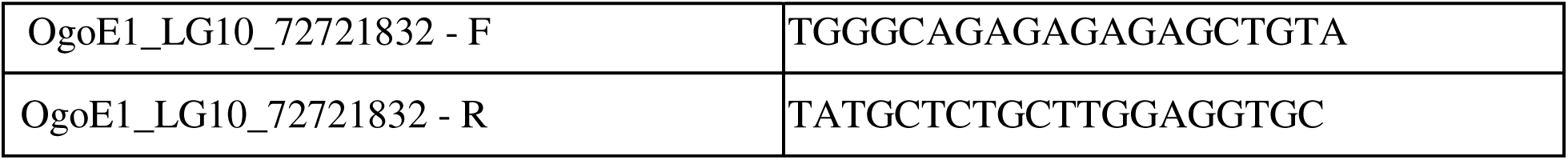

This locus had the strongest association with run timing out of all markers within the *lrrc9* region of the successfully designed primers.

Additional samples were obtained from the 2019 return to the Auke Creek Weir (Table S1) and analyzed for variation at *lrrc9.* Of the 2,094 pink salmon that returned in 2019, 155 individuals were genotyped at OgoE1_LG10_72721832 following the methods of Campbell et al. (*83*). Briefly, dual barcoded sequencing libraries were constructed for the 155 individuals in two 96-well plates (95 samples and one NTC). Individual sample amplicon concentrations were normalized with SequalPrep™ Normalization Plate Kit (Applied Biosystems), libraries were pooled, and then quantified with a Qubit HS kit. The distribution of fragment lengths in pooled libraries was determined with a TapeStation after which molarity was calculated. A double-sided bead size selection was performed using AMPure XP beads (Beckman Coulter) in ratios of 0.5× beads-to-library to remove larger non-target fragments and then 1.2× to retain the desired amplicon. Libraries were sequenced on a MiSeq with 2x75bp chemistry. Paired-end reads for each individual were joined with FLASH2 (*84*) (https://github.com/dstreett/FLASH2). Merged reads were genotyped with the R package GTscore (https://github.com/gjmckinney/GTscore). Individuals that arrived in the first quarter of the run (prior to 23 August) and those that returned in the last quarter of the run (post 28 August) were defined as the early and late component of the return. Genotypic frequencies were compared between the early and late return groups (Fig. S4).

Genotypic frequencies between early and late odd-lineage samples from 2019 had higher divergence than odd lineage samples analyzed with WGS but did not display as high divergence as even lineage samples. In the 2019 early return sample, the vast majority (89%) were early homozygotes and heterozygotes, while the late sample was composed entirely of late-allele homozygotes and heterozygotes in similar frequencies (Fig. S4). The odd WGS dataset included a far higher proportion of late allele homozygotes in the early run sample compared to the 2019 samples, while the odd WGS late sample had allele frequencies similar to the 2019 late sample (Figs. S3C and S4). Divergence between early and late samples was higher in the even lineage than in either dataset from the odd lineage. These results indicate that *lrrc9* influences run timing in both the even- and odd-lineage pink salmon in Auke Creek, but future research is necessary to understand how frequencies vary within and among years.

### Association of *lrrc9* early and late allele frequency in pink salmon populations across the North Pacific Rim

To explore the strength of association of *lrrc9* and run timing in pink salmon, we compared the allele frequency of the early allele among populations across their range and those that spawn between July and October. A total of 1175 pink salmon from 21 collections extending from Russia across the Bering Strait and south though Southeast Alaska were genotyped at the run-timing locus, OgoE1_LG10_72721832 (Table S9). Each of these collections were sampled on a single day, near their peak run timing. The even lineage Duck River collections, in Prince William Sound Alaska, represented both early and late runs into the Duck River and were separated by over a month. The odd-lineage Duck River collection was sampled at a single point intermediate in timing as compared to that of the even lineage. Simple linear regression was used to test for a relationship between Julian date and early-allele frequency. Julian date explained 79.6% of the variation in early allele frequency (F(2,18) = 39.99, *p* < 0.001) and no effect of lineage (*p* = 0.773).

---

**Figure S1.**
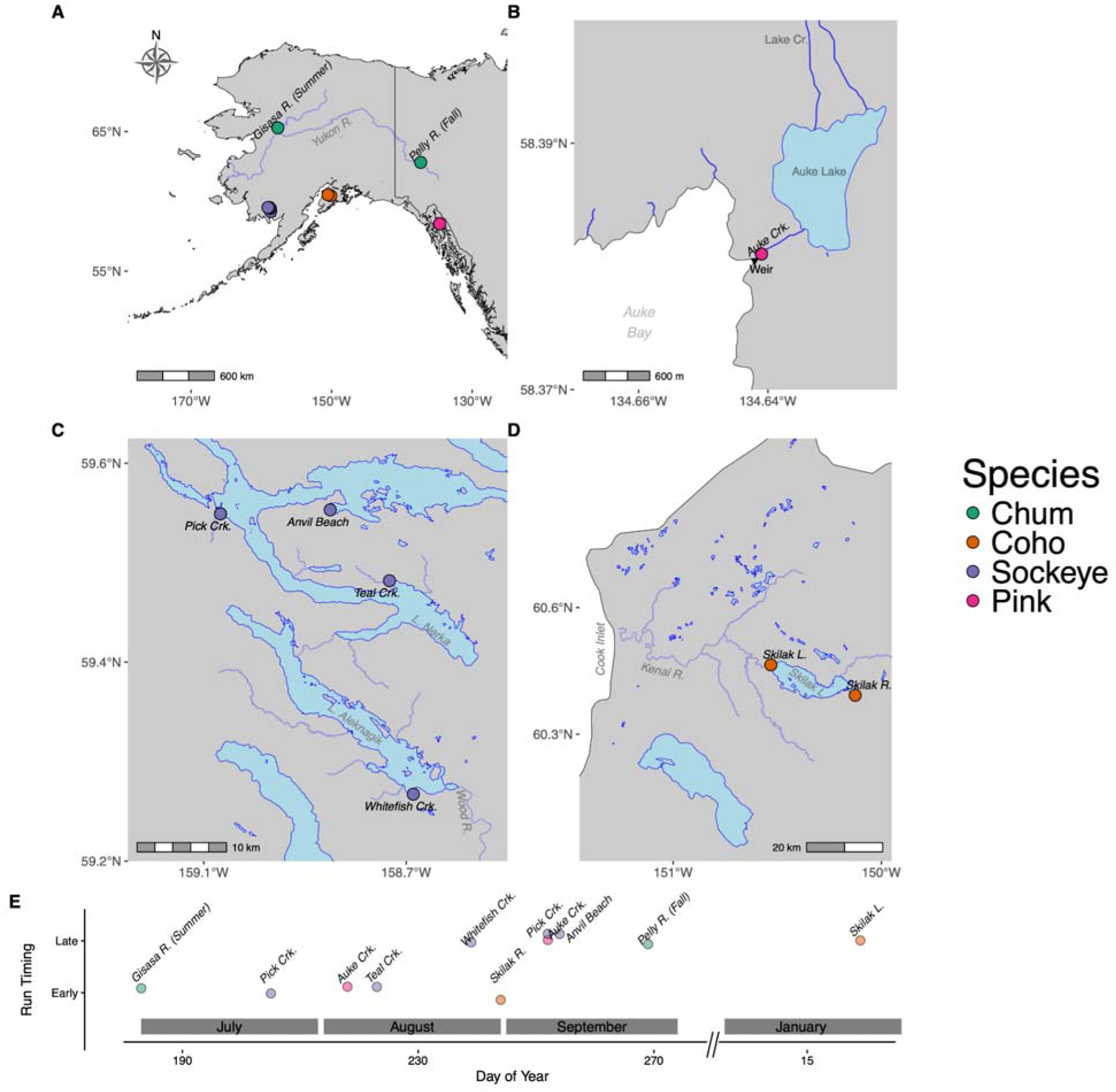
**Geographic and temporal scales over which salmon populations were sampled for whole genome sequencing**. Across Alaska, thirteen collections across Alaska which return to spawn from early July to late January, were included in the study. Map of sampling locations for (A) all species with chum [green] labelled, (B) pink, (C) sockeye, and (D) coho salmon and (E) each population’s estimated peak run timing. Even and odd year lineages of pink salmon have similar peak run timing.

**Figure S2.**
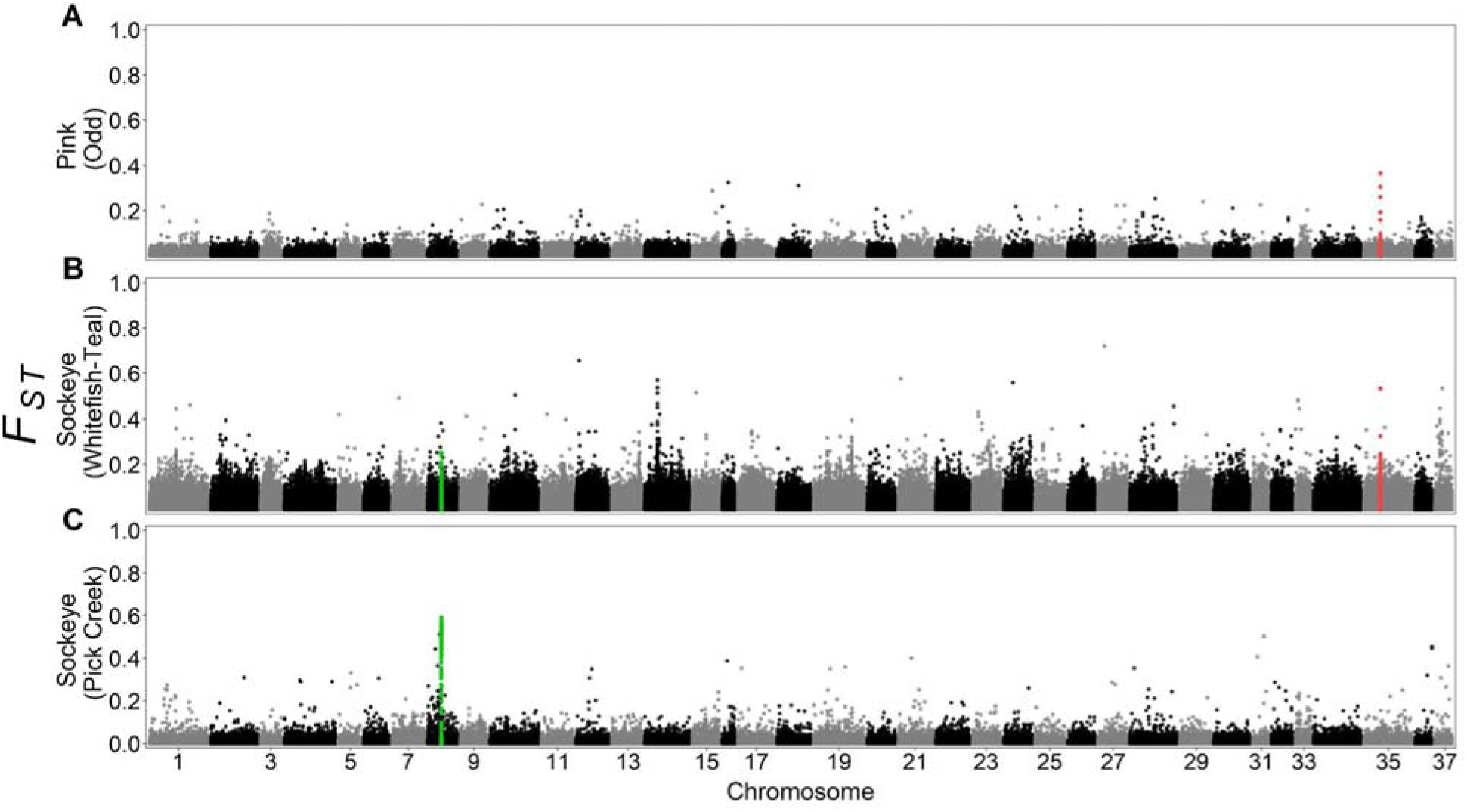
**Manhattan plots of *F*_ST_ between run timing phenotypes across the salmon genome**. Early- and late-run salmon show a weak signal of genetic divergence (*F*_ST_) on chromosome 35 at *lrrc9* (red peak) between (A) odd-lineage pink salmon from Auke Creek and (B) sockeye salmon from Teal Creek and Whitefish Creek, which have an additional weak signal on chromosome 8 at *esr1* (green peak). (C) Sockeye salmon from Pick Creek were highly differentiated at *esr1*, but not *lrrc9*.

**Figure S3.**
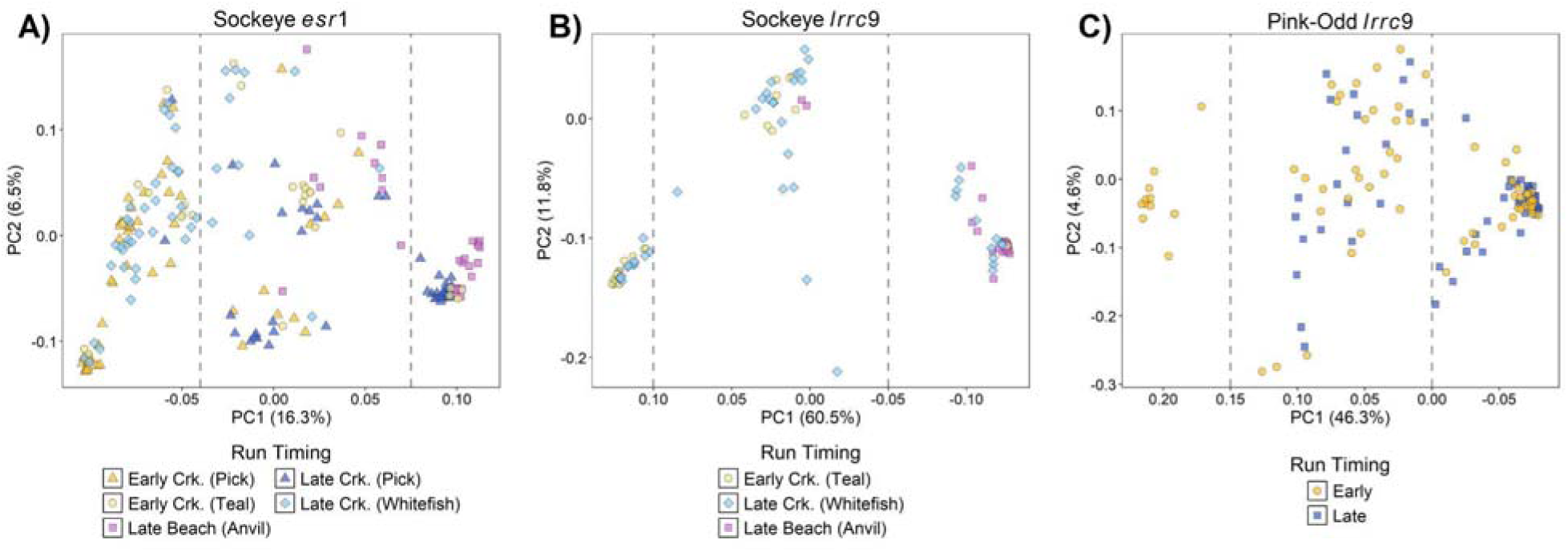
**Genotypic divergence among run timing phenotypes for additional populations of odd-lineage pink salmon and sockeye salmon**. Local PCAs within the *esr1* gene boundary (NC_068428.1:23264160-23645630) in (A) sockeye salmon including Whitefish Creek (late stream) and early and late Pick Creek samples, and the *lrrc9* gene boundary (NC_068455.1:28128954-28169980) (B) sockeye salmon including Whitefish Creek (late stream), and (C) odd-lineage pink salmon from Auke Creek. Sockeye sampling locations are noted in parentheses. Colors correspond to phenotype, and gray dashed lines delineate the designated cutoff for early homozygous (EE, left), heterozygous (EL, middle), and late homozygous (LL, right) genotypes.

**Figure S4.**
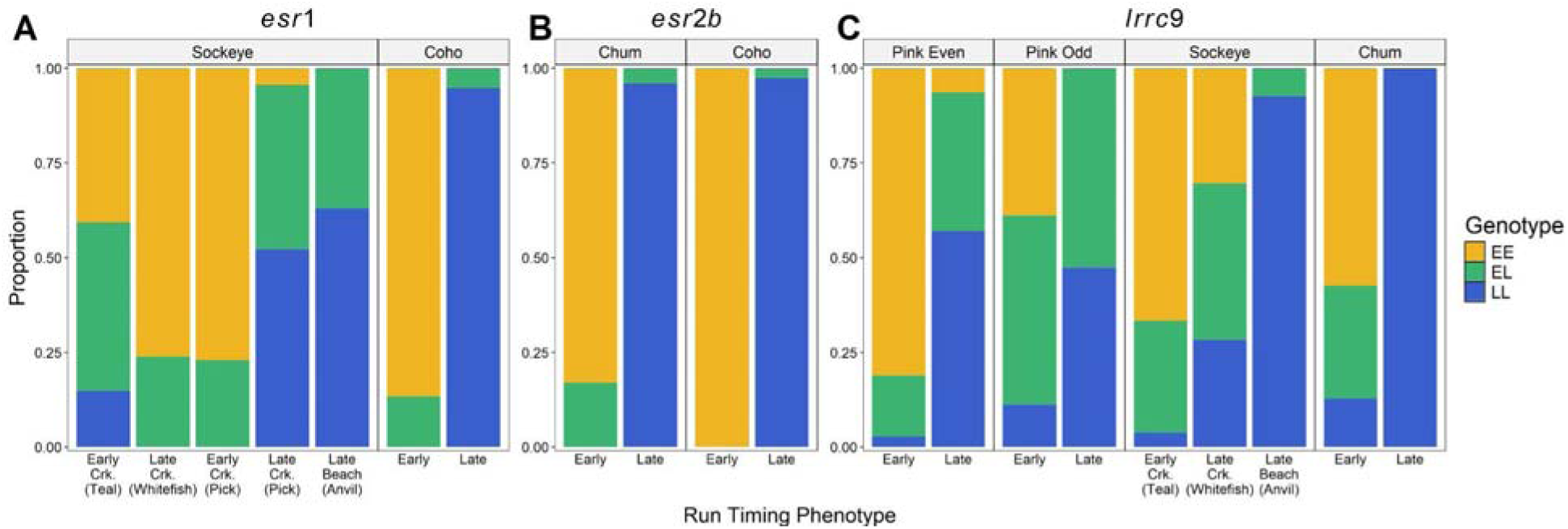
**Genotype frequencies by run timing phenotype and salmon species at three large-effect loci**. Stacked barplots of the proportion of genotypes for each species with a run timing association within (A) *esr1* (B) *esr2b*, and (C) *lrrc9*. The colors indicate genotypes whereas the bars denote the run timing phenotypes within each species. Sockeye salmon collection locations are noted in parentheses. The odd-lineage pink salmon were genotyped with a high-throughput assay marker within *lrrc9* whereas the other populations and species were assigned genotypes from clustering patterns across PC1 within local PCAs at *esr1, esr2b*, and *lrrc9* (see Figures 3 and S3).

**Figure S5.**
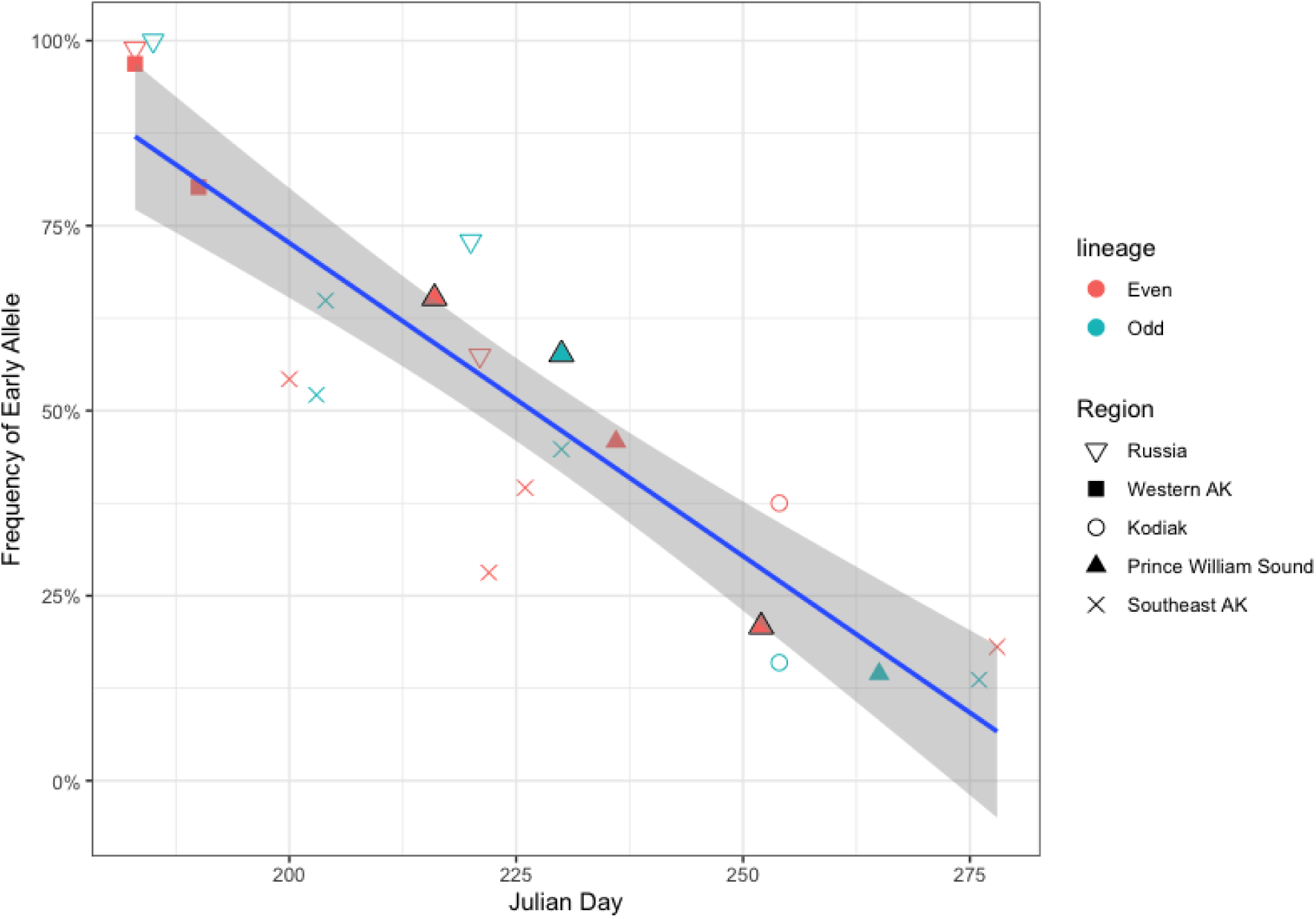
***lrrc9* early allele frequency from pink salmon collections across the north Pacific Rim**. There is a strong negative linear relationship between the early allele frequency and return timing across five regions (r^2^=0.796). Collections from Duck River in Prince William Sound are outlined in black showing variation in early allele frequency in run timing groups within a single return year (even lineage) but maintaining the linear relationship among years.

**Figure S6.**
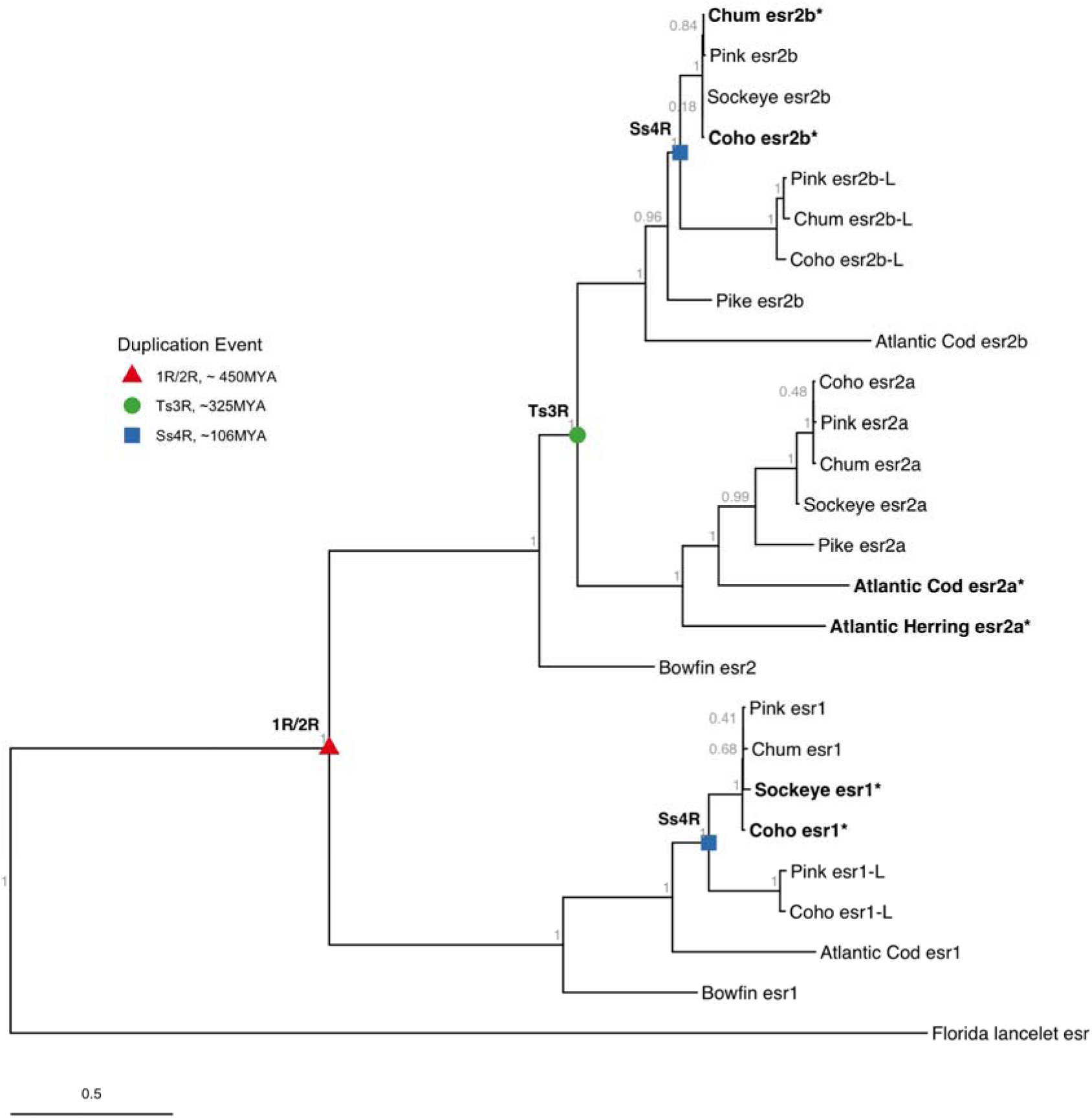
**Estrogen receptor protein evolution through whole genome duplication**. Maximum likelihood tree of estrogen receptor genes. In vertebrates, there are two major branches of er genes, estrogen receptor 1 (*esr1*, *er1*, previously erα) and estrogen receptor 2 (*esr2*, *er2* previously *er*β) which originated after one of the first two whole genome duplications. Within fish, the esr2 gene subtype diverged into two subtypes (*esr2a* and *esr2b*) after the teleost specific whole genome duplication (Ts3R). Salmonid fish display two types of *esr2b* (*esr2b* and *esr2b-L*) and esr1 (*esr1* and *esr1-L*) which arose from the salmon-specific whole genome duplication (Ss4R). Branch lengths represent expected amino acid substitutions per site.

**Figure S7:**
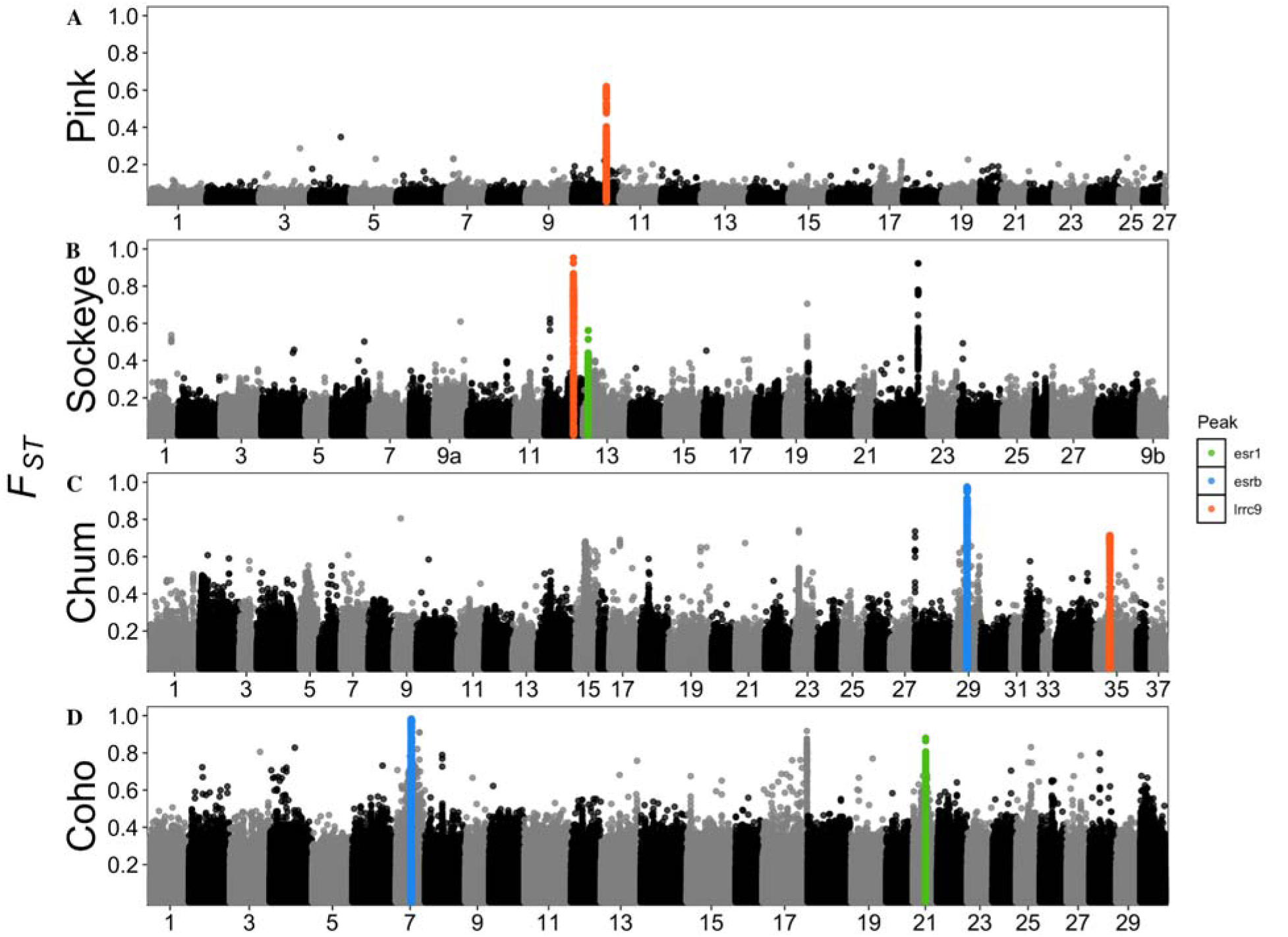
Manhattan plot of genetic divergence (*F*_ST_) between early and late migrating fish in four species of salmon aligned to each species’ respective reference genome. Genome-wide patterns of variability were consistent between species-specific alignments and those to the chum reference genome. Each major effect locus (*esr1*, *esr2b*, and *lrrc9*) from duplicated genomic regions were resolved. Additional peaks on chromosome 22 of sockeye (*ophn1*) and chromosomes 15 (*clcn2-L*) were not shared among species and may show elevated *F*_ST_ due to differences in spawning habit rather than migration timing. The peak on chromosome 28 (*thyrotropin*) of chum and chromosome 17 of coho salmon (*ngfa*) are in similar genomic regions, but the peak on coho was not significant when aligned to the chum salmon reference.

